# 25 Years of Molecular Biology Databases: A Study of Proliferation, Impact, and Maintenance

**DOI:** 10.1101/279067

**Authors:** Heidi J. Imker

**Affiliations:** University Library, University of Illinois at Urbana-Champaign

**Keywords:** databases, research infrastructure, sustainability, data sharing, molecular biology, bioinformatics, bibliometrics

## Abstract

Online resources enable unfettered access to and analysis of scientific data and are considered crucial for the advancement of modern science. Despite the clear power of online data resources, including web-available databases, proliferation can be problematic due to challenges in sustainability and long-term persistence. As areas of research become increasingly dependent on access to collections of data, an understanding of the scientific community’s capacity to develop and maintain such resources is needed.

The advent of the Internet coincided with expanding adoption of database technologies in the early 1990s, and the molecular biology community was at the forefront of using online databases to broadly disseminate data. The journal *Nucleic Acids Research* has long published articles dedicated to the description of online databases, as either debut or update articles. Snapshots throughout the entire history of online databases can be found in the pages of *Nucleic Acids Research*‘s “Database Issue.” Given the prominence of the Database Issue in the molecular biology and bioinformatics communities and the relative rarity of consistent historical documentation, database articles published in Database Issues provide a particularly unique opportunity for longitudinal analysis.

To take advantage of this opportunity, the study presented here first identifies each unique database described in 3055 *Nucleic Acids Research* Database Issue articles published between 1991-2016 to gather a rich dataset of databases debuted during this time frame, regardless of current availability. In total, 1727 unique databases were identified and associated descriptive statistics were gathered for each, including year debuted in a Database Issue and the number of all associated Database Issue publications and accompanying citation counts. Additionally, each database identified was assessed for current availability through testing of all associated URLs published. Finally, to assess maintenance, database websites were inspected to determine the last recorded update. The resulting work allows for an examination of the overall historical trends, such as the rate of database proliferation and attrition as well as an evaluation of citation metrics and on-going database maintenance.

## INTRODUCTION

In the past 25 years online database technologies, and especially web-available databases, have transformed scientists’ use of research data.^1^ Online resources offer researchers unencumbered access to digital content regardless of who they are, where they are, or when they are working. While easy access is undeniably powerful, the sustainability of proliferating scientific resources comes with many challenges. The cost of computation and storage has decreased steadily for decades and are no longer a rate-limiting factors. However, database providers are still challenged to acquire funds—both initial and ongoing—to cover staff and other technology costs, including efforts to find and retain skilled professionals, update and secure systems, and ensure data are accurate and current. In light of these challenges, the proliferation of online resources and data itself has been closely chased with concerns of sustainability (Ember and Hanisch 2013; Guthrie et al. 2008; Kalumbi and Ellis 1998).

The recent intensified interest in data, including data science, big data, and data analytics, has been coupled with a push by funding agencies, publishers, and scientists themselves for increased access to research data. A memo from the Office of Science and Technology Policy explicitly called for public access to the products of federally funded research, including data (Holdren 2013). Sharing data via web-available databases is a common mechanism, yet this creates a conundrum: how can more access to data be attained when there is already intense competition for funding to maintain current databases? A better understanding of scientific community’s capacity to develop and maintain web-available databases would help our understanding of the sustainability of such resources.

To date, database studies have tended to fall on either end of a spectrum. On one end, studies have covered small or medium sized samples, allowing careful inspection of individual resources (Attwood et al. 2015; Kirlew 2011; Marcial and Hemminger 2010). At the other extreme, studies have evaluated Scientific Data Analysis Resources (SDARs) by proxy through mining tens of thousands of URLs in the academic literature (Wren et al. 2017). Both strategies have provided valuable insights, but studies that have straddled the middle are rarer. This work aims to complement these efforts by gathering a historical sampling of databases that is large enough to evaluate trends in proliferation and impact, but also granular enough to allow identification of sizable subsets of individual databases to assess for maintenance. An overarching goal of this study is to provide the ability to identify subsets of databases for further analysis, both as presented within this study and through subsequent use of the openly released dataset.

Absent of the ability to evaluate all online databases across all academic disciplines, a large, diverse, and well-documented sampling of databases is required. Molecular biology is a far-reaching field where principles and techniques are core to nearly all other disciplines in the life sciences, from immunology to biophysics. Correspondingly, review of molecular biology databases compendia show they cover a wide swath of physical, chemical, biological phenomenon across a range of organisms (Fernández-Suárez and Galperin 2013) and are often created with the intent to be useful to a variety of biologists (Galperin and Cochrane 2009). Additionally, the molecular biology community has progressively created standards, technology, and community norms to support wide-spread reuse of data.

The desire to develop and maintain community resources is not unique to molecular biology; many other disciplines—ranging from astronomy to hydrology to archaeology—have a high appetite for community resources, but as academic endeavors all are similarly constrained by the funding options available. Sustainability of data resources has been a topic of active discussion for many years and continues to be an unresolved issue (OECD 2017). However, the long and enterprising history of database adoption in molecular biology provides a wealth of diverse databases with mature lifecycles to study. Some are under the purview of large, well-established government organizations, such as ArrayExpress from EMBL-EBI, and others are created by small university-based research groups such as RDP: Ribosomal Database Project. Especially as a population, these databases’ lifecycles provide an opportunity to examine the variable conditions under which online databases are created, utilized, and maintained. Given the underlying similarities associated with the challenges of database sustainability across all academic disciplines, lessons gleaned from these molecular biology and bioinformatic resources are likely informative for other domains.

With this in mind, the study presented here aimed to create a census of sufficiently documented molecular biology databases to answer several preliminary research questions and serve as fodder for future work. Namely, the questions addressed herein include: 1) what is the historical rate of database proliferation versus rate of database attrition?, 2) to what extent do citations indicate persistence?, and 3) are databases under active maintenance and does evidence of maintenance likewise correlate to citation? Articles published in the annual *Nucleic Acids Research* (*NAR*) “Database Issues” were used to identify a population of databases for study. *NAR* is an prominent, well-respected journal in the molecular biology and bioinformatics communities that has long served as vehicle for description of data and databases. *NAR* published articles related to compilations of data, especially gene sequences, for many years, but it wasn’t until 1991 that a dedicated supplemental *NAR* issue focused on databases (Fernández-Suárez and Galperin 2013). The first formally named Database Issue was published in 1993, but for the purpose of capturing these earlier efforts, the study here includes these two preceding supplemental issues from 1991 and 1992 to provide a fuller history of database development. This affords an exceptionally long period of time given the pace of technology change and development of science itself.

To make use of this 25 year history, the author mined the articles published within these issues to answer fundamental questions about the databases reported. Subsequently, this work represents the identification of 1727 unique databases and the collection and analysis of descriptive statistics for each, including year debuted in a *NAR* Database Issue, current availability, as well as article and citation counts for all associated *NAR* Database Issue publications for each of unique database. Additionally, to address gaps in our current knowledge, the apparent date of last update was also determined through visual inspection of web pages for all databases currently available. While assessment of URL response provides a binary understanding of database availability, e.g. either “accessible” or “not accessible,” the apparent date of last update is an important but previously unreported metric that provides a window into the activeness of a database’s current operation.

## METHODS

### Dataset Assembly

The initial source of metadata for each database article, including Author, Title, Year, Cited by, and DOI, was gathered from the Scopus abstract and citation database on December 6, 2016. For the purposes of this study, the issues included as “*NAR* Database Issues” include the formal Database Issues from 1993 forward as well as two preceding supplemental issues in 1991 (Vol 19, Issue suppl) and 1992 (Vol 20, Issue suppl). No Database Issue was published in 1995, resulting a total of 25 issues subjected to query.

Only articles that describe a specific resource were included in analysis. Of the 3115 records extracted from Scopus for *NAR* Database Issues, 3055 articles were included. The articles removed include editorials (n = 10) and overall descriptions of resources at the National Center for Biotechnology Information (n = 17) and the European Bioinformatics Institute (n = 7); both of which commonly mention a dozen or more resources, including non-databases. Articles describing generic topics such as Oracle, Wikipedia, legal interoperability, and regular expressions (n = 4) were also removed. Finally, a small number of duplications and other indexing errors (n = 22) were also discovered and removed, leaving a final count of 3055 articles in the sample.

### Identification of Unique Databases

Unique databases published in *NAR* Database Issues were identified from individual articles. In 2009 article titles were standardized to “Database Name: short description,” and database names were readily extracted using this syntax. However, previous years offered no such standardization, and titles required manual review. When a name could not be identified from the title, the abstract was reviewed. If this failed, the articles themselves were reviewed. While the vast majority of articles describe a single resource, a small portion (n = 37) were found to describe two or more resources. Assuming equal representation, citations for these articles were divided by the number of resources within to provide a citation count for each database described.

Articles fall into two categories: “debut” articles which cover a database’s first appearance in an *NAR* Database Issue and “update” articles which provide an update of a database previously described in an *NAR* Database Issue. A “debut” year is not necessarily a creation year since a database may have existed for several years prior to publication. While annual updates were common in early years, submission of update articles were later limited to every other year, with a few acceptations for major resources such as GenBank, DDBJ: DNA Data Bank of Japan, and ENA: European Nucleotide Archive (Galperin and Cochrane 2009). With the expectation that a given database may be associated with multiple articles over the years, the next step was to determine how many articles describe each unique database. This enables a fuller understanding of a database’s history through determination of metrics over multiple years. In order to accomplish this, extracted database names were checked for standardization and then cross-checked to ensure that a single unique database was not misidentified as two (or more) separate databases through cryptic reference in separate articles. Confounding issues complicate this task such as the names of databases changing over the years or mergers between resources. A set of criteria were defined to guide name standardization (Imker 2018), and for additional validation URLs and author names were also checked to identify potentially missed matches.

### Database Availability and Updating

Only web-available databases (n = 1714) were assessed for availability. Database URLs were recorded from article abstracts and tested manually once between the period of Dec 19, 2016 – Feb 22, 2017 to inspect evidence of ongoing updating and maintenance. For databases with more than one URL, either as a result of multiple articles or multiple access points (e.g. mirror sites), only one functional URL had to direct to the database in order for the database to be categorized as available for this study. In some cases, URLs were not functional and threw typical client or server errors. However, in other cases the URL was functional, but did not provide access to the database. Examples include webpages that contain a discontinued notice or a redirect either to a related, but generic website (e.g. the home page of a university) or to an entirely unrelated website (e.g. an e-commerce site). In these cases, although these URLs technically resolve (e.g. return the HTTP status message 200) these were recorded as the database being unavailable if no other URLs resolved to the actual database.

For URLs that did resolve appropriately, the websites were inspected to determine an apparent date of last update as a way to infer if the database was under active maintenance. Date stamps for page updates were not counted since these can be auto-generated every time a webpage is accessed. Likewise, copyright dates in footers can be dynamically updated and were not counted as evidence of active maintenance. The home page was scanned, and any areas or navigational elements labeled as “announcements,” “news,” “updates,” “versions,” or “releases” were inspected and the year of the update was recorded, if found. A database for which no date could be located does not necessarily indicate the database is not being maintained, simply that the date of last update could not be determined. From this analysis an date was located for a total of 591 databases covering the entire 1991-2016 time frame.

### Validation and Assignment of Identifiers

Identifiers were assigned to standardized database names to facilitate analysis. As another check of accuracy and also to enhance future usability of the dataset, identified databases were mapped to Molecular Biology Database Collection (MBDC) IDs and assigned the same identifier, where present. A list of MBDC IDs and associated database names were collected from the Oxford University Press website on December 8, 2016. Databases within the MBDC were matched with the standardized database names assigned in this study; when no match was recognized or resources were named ambiguously, the MBDC page was accessed to determine the associated article. As expected, not all MBDC databases correspond to articles published within in *NAR* Database Issues since the MBDC accepts database submissions without an associated *NAR* publication. Likewise, not all databases identified in this study are represented in the MBDC since the MBDC is meant to represent a current list of active databases and defunct databases are culled from the collection on an annual basis. A total of 322 databases could not be mapped to MBDC ID, leaving 1405 that map to the MBDC. The 322 that could not be mapped were given an identifier unique to this study.

### Assignment to Citation Quartiles

Two major issues that can skew citation analysis include 1) the number of article citations strongly correlates with time since publication resulting in disproportionately low citation counts for new articles and 2) raw citation counts fluctuate wildly and are difficult to compare side-by-side. To address the first issue, publications from years 2013-2016 where excluded from citation analysis based on deviation from average citations/article seen over the preceding period (see Supplemental Figure S1). To address the second issue, percent rankings were adopted as a method to normalize citations (Waltman and Schreiber 2013). In this analysis, each article’s ranked placement within an issue (“Issue Percent Rank”) was calculated by ordering articles by number of citations and determining each article’s percentile. For databases with a single debut article published prior to 2013 (n = 842), this represents the database’s overall Percent Ranking. For databases with a debut article plus additional subsequent update articles published prior to 2013 (n = 515), the Issue Percent Rank was averaged across all articles to determine the database’s overall Percent Ranking. With this calculated, each database was binned into its corresponding Citation Quartile; e.g. databases with a Percent Ranking between 1.0 and 0.75 assigned to the first quartile, and so forth.

### Data Analysis

The dataset was reshaped and analyzed to obtain aggregate issue-level, article-level, and database-level metrics using base functions in the statistical programming language R as well as the packages tidyverse (Wickham 2017b) and stringr (Wickham 2017a). The packages gglot2 (Wickham 2009) and ggpmisc (Aphalo 2016) were used to visualize results using the Tol technical specifications for color schemes (Tol 2012). To assess potential statistical significance of increasing or decreasing trends for quartiles, a chi-squared test for trends in proportions was used (Dalgaard 2008). To assess the distribution within quartiles Pearson’s chi-squared test for count data was used (Agresti 2007). Data for trend lines were selected to maximize linearity as determined by calculating the second derivative to identify data points associated with maximum curvature. The R scripts written are available with the associated dataset (Imker 2018).

## RESULTS AND DISCUSSION

### Growth for NAR Database Articles and Debuted Databases

Although *NAR* occasionally published small, supplemental issues devoted to compilations of gene sequences throughout the 1980s, articles in a supplemental issue from 1991 began to show a marked movement towards the adoption of database technologies. Both the 1991 and 1992 issues contained many articles that described databases that already existed online or described precursor efforts to develop future web-available databases. The first supplemental issue in 1991 contained just 18 articles and represents a snapshot of change. Several of the databases described were only available via postal delivery of physical media such as floppy disks, CD-ROM, or even paper printout (e.g. see Wells and Brown 1991; Wada et al. 1991; Gupta and Reddy 1991).

Others offered server access through Gopher or FTP, and yet others indicated an expectation to transition to new forms of access, such as “new-style databases” offered via centralized services (e.g. Giannelli et al. 1991). The first databases accessible via the World Wide Web appeared in 1994, and adoption quickly spread. In told, 53 databases published in *NAR* Databases Issues started either prior to or concurrent with the advent of the World Wide Web. Of these 53 databases, 40 (75%) transitioned to a web-accessible format and 18 (45%) of those remain available as December 2016 (Imker 2018). The last publication that did not offer web access, for the WT1 Gene Database, occurred in 1998 (Jeanpierre et al. 1998). The fact that entire issues, which had grown 5-fold to over 100 articles by 1999, could exclusively cover web-accessible databases is a remarkable change in just five years.

By 1999 such a large number of databases had been published that the Database Issue needed a resource itself. First compiled as a list and then moved online a year later, the Molecular Biology Database Collection (MBDC) serves as a continually updated auxiliary resource that facilitates researchers’ ability to locate current, relevant databases (Baxevanis 2000). As submissions to the *NAR* Database Issue continued to balloon through the early 2000s, a 2009 Database Issue editorial reiterated strict publication criteria that emphasized “thoroughly curated databases that are expected to be of interest to a wide variety of biologists, primarily bench scientists” with a preference towards unique, web-accessible, and persistent databases not described elsewhere (Galperin and Cochrane 2009). In 2010 it was announced that a new journal, *Database: The Journal of Biological Databases and Curation*, was launched by Oxford University Press with the hope that this new journal, along with *Bioinformatics*, would provide an appropriate venue for databases deemed not appropriate for inclusion within an *NAR* Database Issue. These changes checked growth, and in the following years the issue size leveled out to an average of 183 articles published per year over 2011-2016 (Figure 1).

**Figure 1.**
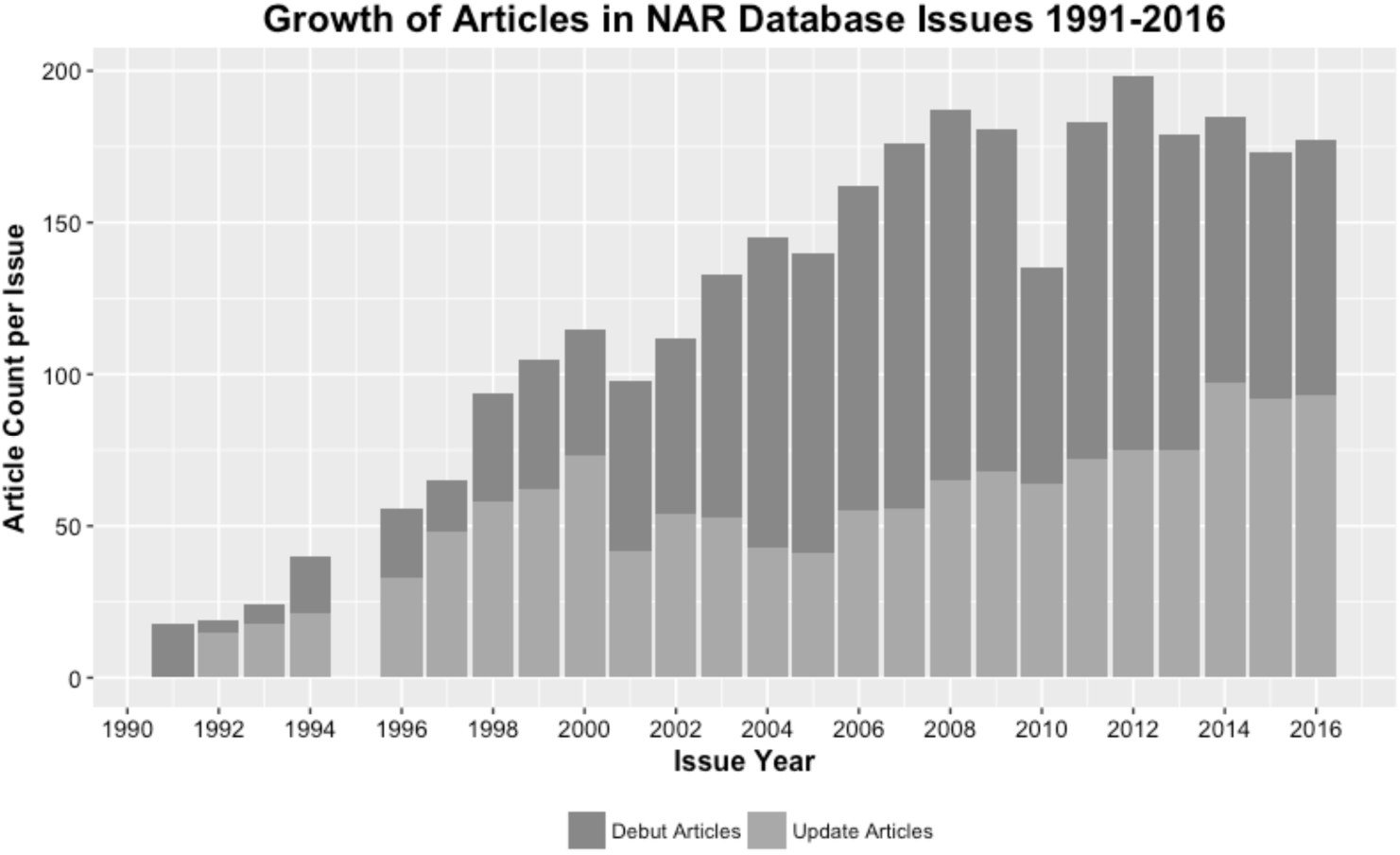
*NAR* Database Issue growth between 1991-2016. Stacked bars represent the total number of articles published in a given year and are further broken down into debut articles (dark grey) and update articles (light grey). Note that here (and figures throughout), the absence of data for 1995 is the result of no publication of a Database Issue that year.

As dissemination of data via the web became the norm, the number of unique databases debuted began to accumulate. Debuts grew steadily throughout the 1990s with a sharp inflection around the turn of the century (Figure 2). The data gathered here show that for over the last decade the rate of database debut has closely followed a linear trend. This rate is necessarily limited due to the capacity of the *NAR* and the community to handle editorial and peer review responsibilities for associated articles. Despite this constraint, a brisk “proliferation rate” of 104 database debuts/year can be calculated for 2002 – 2016.

**Figure 2.**
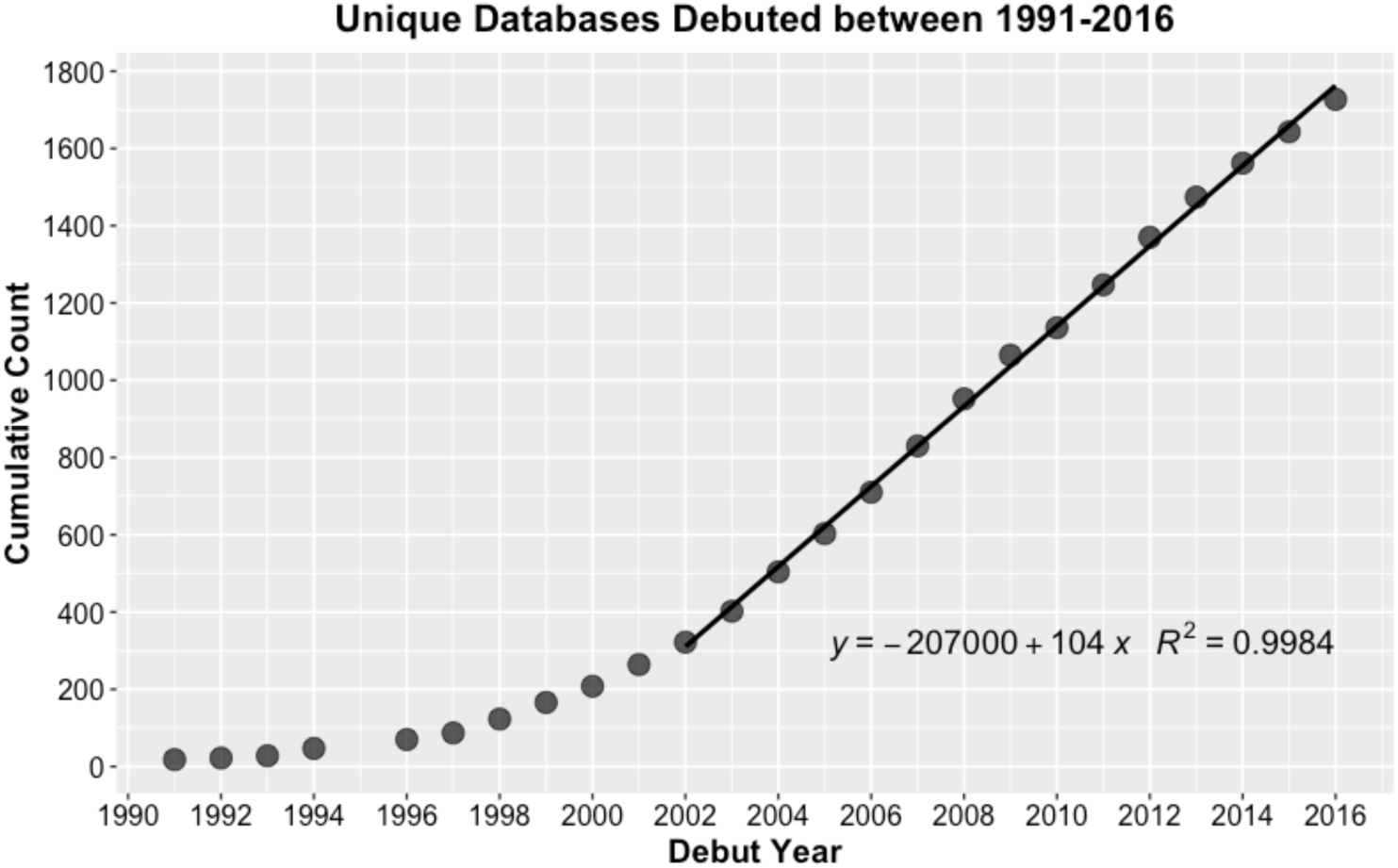
The accumulation of unique databases debuted in *NAR* Database Issues between 1991-2016 shows a continual trend of increased growth overall and linear proliferation in recent years, with maximum curvature in 2002.

*NAR* Database Issues focus on highly-developed, well-curated databases that are expected to be of broad interest to research communities. More databases are developed that are not included within *NAR* Databases Issues, although a database provider survey conducted for ELIXIR in 2009 indicated that *NAR* dominates database publication (Southan and Cameron 2017). Thus the work here is expected to provide a robust sampling, but also represents a “lower limit” of database proliferation. Especially when considered as a lower limit, this proliferation rate illuminates the magnitude of the sustainability issue. An annual growth rate of 104 new, unique, and high-quality databases that are expected to persist well into the future is not a trivial number. Given erratic funding for scientific research, the ability to indefinitely sustain a growing number of resources—not matter how high of quality—is a major cause for concern (Baker 2012; Merali and Giles 2005). Indeed, multiple successful resources published within *NAR* Database Issues have suffered. Example databases within this dataset that have amassed >1500 total citations and yet have moved to variable subscription or donation models include OMIM: Online Mendelian Inheritance in Man, TAIR: The Arabidopsis Information Resource, TRANSFAC, and even KEGG: Kyoto Encyclopedia of Genes and Genomes. As a database with 16,832 total citations, the second highest found in this study, KEGG’s value to the community is not ambiguous. Thus the proliferation rate found for databases published in *NAR* Database Issues puts pervasive funding issues in starker light as database creators, funders, and researchers grapple with sustainability.

### Measures of Database Impact

Databases require financial and organizational support not only for initial creation, but for on-going maintenance and improvement. Most community databases, like other research infrastructure resources, function on unstable grant and institutional support (Bastow and Leonelli 2010; Kalumbi and Ellis 1998). This is despite their critical role in modern science; for example, Wren found bioinformatic resources were disproportionately overrepresented among articles with the most citations between 1994-2013 (Wren 2016). Because of the lack of stable support, evidence of impact is highly sought after in order to prioritize allocation of funds and provide compelling justifications for continual government or institutional backing (Mayernik et al. 2017).

Citation analysis has well-acknowledged limitations, yet remains a common method to assess impact in many areas of academia. Critics note that citations are not comprehensive indicators of impact since authors rarely cite everything that could or should be attributed (MacRoberts and MacRoberts 2017). This is true for databases as well, and studies examining full-text journal articles found data citation is often under-reported (Jonkers et al. 2014; Mooney 2011). Two issues are especially confounding. Without a formal publication, databases in and of themselves generally do not map to a standard bibliographic format required by publishers. Additionally, researchers themselves often do not recognize their data sources as citable. Consequently, references are often relegated to in-line text or not mentioned at all (Mayo, et al. 2016). Jonkers et al. found that citations for HAMAP and SWISS-2DPAGE, two databases included in this work, were underrepresented by 11.1 % and 27.8 %, respectively, when in-text mentions were evaluated and results were even more variable when other ExPASY resources where examined (Jonkers et al. 2014).

In light of this underrepresentation, a major push to enable formal citation of data is underway. The Joint Declaration of Data Citation Principles brought together a group of interested parties to develop a set of principles to encourage data citation and acknowledge “data should be considered legitimate, citable products of research” (Martone 2014). The organization DataCite, founded in 2009, issues digital object identifiers (DOIs) as a way to assign persistent identifiers to datasets (Neumann et al. 2014). According to DataCite’s statistics dashboard, by the end of 2017 DataCite issued over 12 million DOIs.^2^ Additionally, large publishers such as Elsevier now include explicit encouragement in author guidelines to cite the data that support articles. PLOS not only has a strict availability requirement for data that underpins articles, but likewise includes guidance on how to cite databases and repositories. The momentum towards greater acknowledgement of data and data resources is strong, and citation is clearly expected to be an essential metric.

The idea of data citation is certainly not new to database providers, and statements requesting citation can be found within early *NAR* Database Issue publications (e.g. Jeanpierre et al. 1998). Although the reinvigorated emphasis is welcome, change continues to percolate slowly through academic communities (Mayo et al. 2016). In the meantime, *NAR* Database Issues publications offer an opportunity to study data citation by proxy and also follow accepted criteria in citation analysis; i.e., citations evaluated in aggregate, and through comparisons within the same year, same type of publication, and from the same discipline (Leydesdorff et al. 2016).

As of December of 2016, citations for all 3055 Database Issue articles included in this analysis totaled 385,235. New issues show an expected citation lag, but cumulative citations show linear growth with each new issue at a rate of 22,800 citations/issue/year between years 1999-2012 (see Supplemental Figure S1).

The results in Figure 3 show database availability correlates to number of citations. These results corroborate a recent analysis of SDARs, which also included databases (Wren et al. 2017). However, this relationship does *not* hold true for recent articles since citation lag is readily apparent for recent publications. As such, these results indicate that a low number of citations within a few years of debut cannot be used to anticipate that a given database will not be popular. In their survey of >200 database providers, Southan and Camerson found over 90% of providers had less than 3 years of secured funding, and other surveys have indicated similar instability (Attwood et al. 2015). Since it is likely that additional support will be required during the first few years of a database’s debut, early citation metrics should not be relied on as a raw measure of impact. This is an especially important point for those submitting (or evaluating) funding proposals.

**Figure 3.**
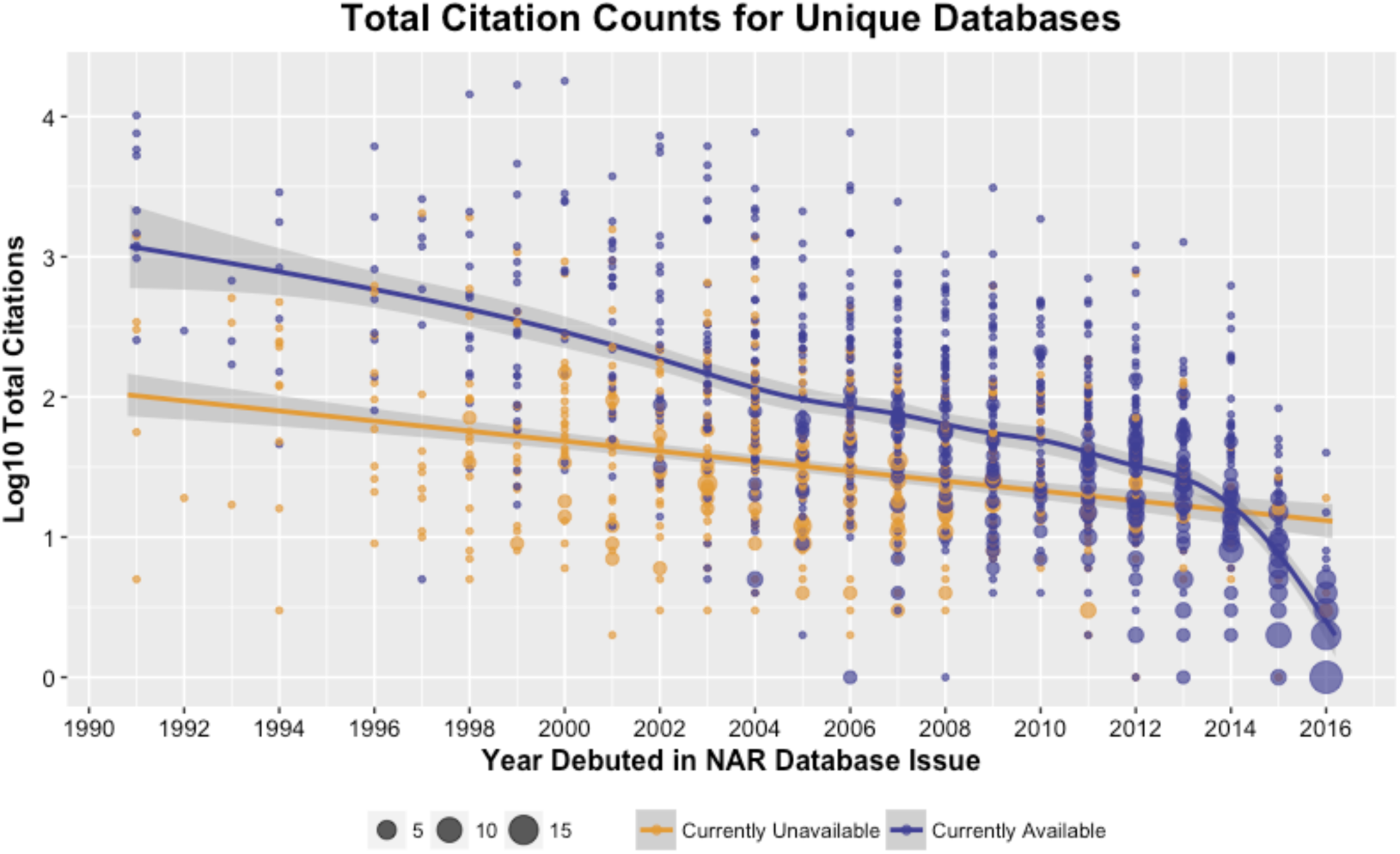
Total citations (displayed as log10) across all articles that describe a given database are higher for databases that remain available (purple) than for those no longer available (orange). In this analysis of raw citation counts, the time dependence of citations is stronger for currently available databases than those that tested as currently unavailable. Points are scaled to represent the number of databases for a given citation count and debut year. To better fit local means, especially for more recent years, a generalized additive model function was used, and curves are shaded with 95% confidence intervals.

Citations are known to vary wildly, as is true in this analysis. For example, the 2000 article for RCSB PDB: Protein Data Bank is a notable outlier with 16,498 citations. This is 3 times the second most cited article which has 5558 citations for KEGG: Kyoto Encyclopedia of Genes and Genomes, also published in 2000. To mitigate this issue of skewing and enable more robust comparison across years, article citations were normalized within issues to calculate a Percent Rank which then was averaged across all *NAR* Database publications for a given database. To account for citation lag only articles and associated databases from 1991-2012 are included in this analysis. Percent Rank was then used to bin databases into Citation Quartiles for analysis (Table 1).

**Table 1.** Citation Analysis for Databases with Articles Published 1991-2012

| **Citation Quartile** | **Databases Count (% Total)** | **Articles per Database Average (STD)** | **Raw Citations Sum (% Total)** | **Currently Available Count (% w/in Quartile)** |
| --- | --- | --- | --- | --- |
| **1** | 193 (0.14) | 3.3 (3.3) | 268,704 (0.72) | 155 (0.80) |
| **2** | 338 (0.25) | 2.4 (2.8) | 72,889 (0.20) | 232 (0.69) |
| **3** | 385 (0.29) | 1.7 (1.6) | 23,528 (0.06) | 236 (0.61) |
| **4** | 440 (0.32) | 1.3 (0.90) | 7,211 (0.02) | 193 (0.44) |
| **Totals** | 1,356 | -- | 372,332 | 816 |

The data show a small proportion of databases in the 1^st^ quartile (14%) ultimately represent the majority of the *NAR* Database Issue citations (72%). For some databases, these citations may accumulate over multiple articles. This percentage jumps to 92% when the top two quartiles, representing 39% of the set, are considered. The percentage of databases still available significantly decreases as the bottom quartile is approached (𝒳^2^ = 87.508, *p* < 2.2e-16). In availability testing for this analysis, 80% of the databases in the 1st quartile remain accessible. Considering the apparent impact of databases in this quartile, the absence of 20% is potentially problematic. However, it is possible that these databases migrated in a manner that was not detected in this analysis or have simply outlived their usefulness. An in-death analysis of these absent, high-ranking databases and the consequence of their absence on the community would aid this assessment.

On the other end of the spectrum, despite the relative lack of citations for databases in the 3^rd^ and 4^th^ quartiles, URL testing showed that 61% and 44%, respectively, persist. Although one might be concerned about the loss of any resource, this may be a natural consequence if low citations indicate the databases were not of as great of utility as originally expected. For example, Galperin suggested some databases may be duplicative or suffer from a narrow scope (Galperin 2006). In that light, the retention of over 400 databases within these two quartiles is noteworthy. Closer inspection of these resources will help us understand what drives persistence with little external validation in the form of citation. Indeed, there may be important lessons to learn from these databases to apply more broadly. For example, is persistence achievable because of easy to maintain systems? A personal commitment? A different (and possibly more appropriate) measure of impact? With additional validation and refinement, assignment of citation quartiles provides a framework for targeting subsets of databases to assess in more detail, for example, to assess for support needed to sustain, sunset, or even resurrect resources.

### Persistence and Maintenance of Databases

Databases published in *NAR* Database Issues are expected to persist. If databases are neglected, the *NAR* Database Issue editorial from 2009 stated “respective senior authors (and in some cases, their host institutions) will be prevented from publishing new papers in the *NAR* Database Issue” (Galperin and Cochrane 2009). Indeed, these expectations appear to be effective since the rate at which databases published in *NAR* Database Issues become defunct is low compared to other measures. For example, a 2014 study found that positive responses to requests for data supporting published articles fell by 17% per year (Vines et al. 2014). In contrast, the availability of the databases tested here decreased by only 3.8% per year from 2001-2016 (Figure 4), which corroborates an earlier estimation of < 5% per year for databases included in the MBDC (Fernández-Suárez and Galperin 2013).

**Figure 4.**
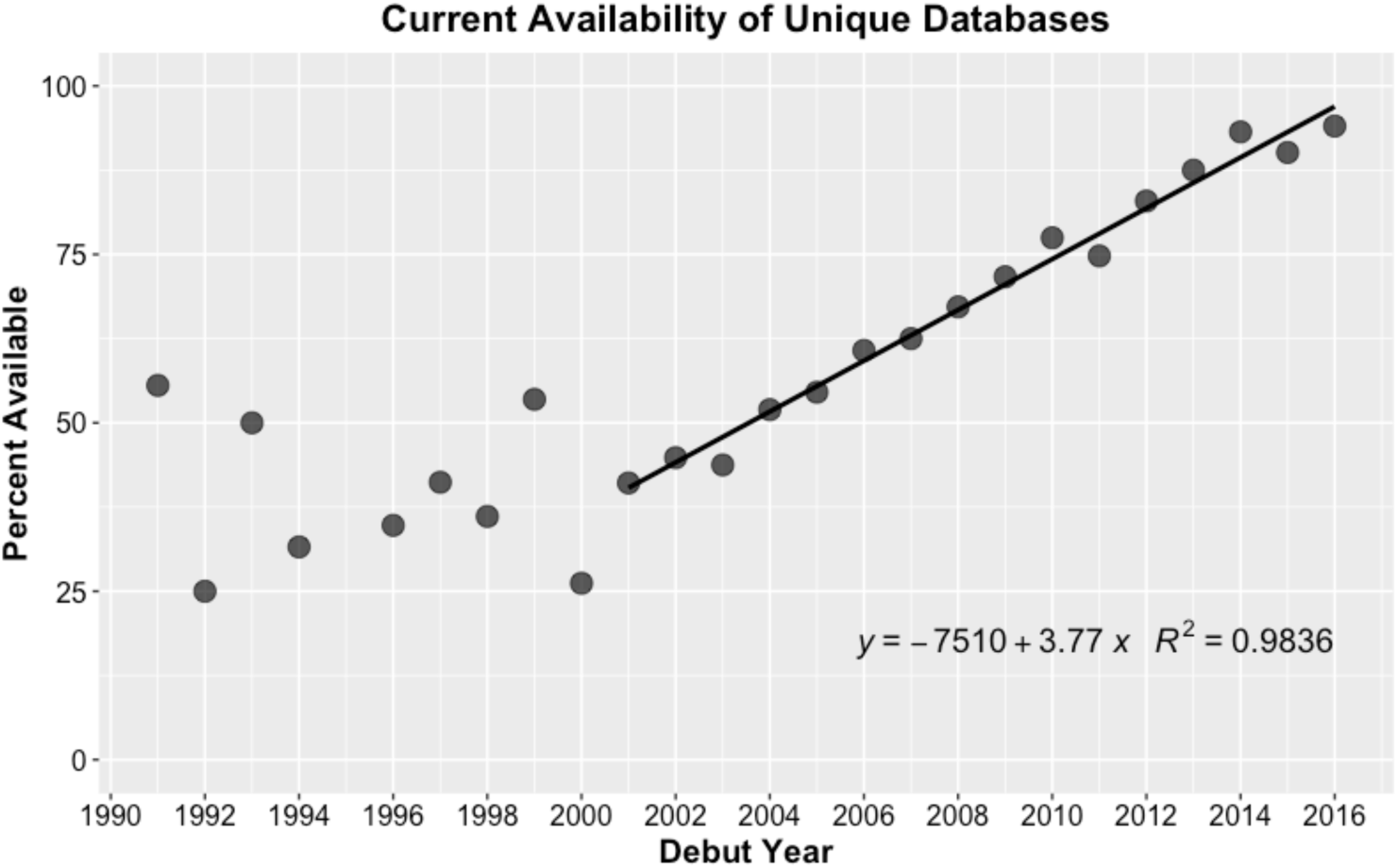
In this study, URLs for each unique database were tested to determine the rate of database attrition. Databases show linear attrition for those debuted between 2001-2016, with maximum curvature in 2001. For earlier years, although the data here indicate retention is highly variable, database availability ceases to decline steadily.

Interestingly, attrition appears to level off for databases debuted between 1991-2001. This finding is notable as it allows us to quantify the number of databases that are, in fact, persistent for a long period of time (e.g. ≥ 15 years). The current availability of databases that debuted between 1991-2001 averages to 39.5% and equates to 105 databases. One interpretation of this result is that the community is able to sustain a net positive growth in the number of databases that can be supported long term. In this scenario, although the community is unable to sustain all debuts such that substantial attrition of new databases does occur, a portion of new databases are retained long-term and the total population grows. However, an alternative interpretation of this result is that leveling off represents the approximate maximum capacity the community can sustain, such that existing “established” databases must cycle off to make room for newer databases to hold the population steady at ∼100 databases maintained overall. These interpretations represent two very different scenarios, and it is critical that this trend is watched closely in the coming years. Additionally, examination of the 105 long-lived databases in more detail will provide valuable insights into the characteristics that contribute to longevity.

Whether or not a URL continues to resolve to the database is helpful to evaluate if a database is still accessible. However, to measure on-going maintenance of the databases identified in this study, each available database website was inspected for an “update date.” This date may be associated with any number of improvements, such as the addition of new data, development of new features, creation of tutorials, changes to the interface, server upgrades, etc. For the purposes of this study, any such activity was considered an update and was interpreted as evidence that the database is undergoing active monitoring and care. This evaluation is of interest because it signifies on-going effort and helps to assess the need for committed funding.

The same set of currently available databases with articles published between 1991-2012 (n = 816) assigned to citation quartiles was used to analyze maintenance. This subset provided the ability to examine maintenance relative to citation quartiles but also ensured the databases included were old enough that the need for update is likely. The results show that an update date could be located for 434 (53%) of these databases (Table 2). Since the inability to locate an update date could be the result interface complexity or simply that updates are not publicized, the remaining 47% cannot be interpreted as not updated, simply that update is unknown. However, the higher the citation quartile, the more likely that updates were documented, with dates almost twice as likely to be found for a database within the 1^st^ quartile versus the 4^th^ quartile (𝒳_2_ = 46.084, *p* = 1.133e-11). That the presence of an update date, regardless of what the actual date is, correlates to citation ranking is a surprising result and may hint at underlying issues with the usability of the database websites. Readily providing update information allows users to assess how current the database is and may influence users’ interpretation of reliability. Biological databases and other bioinformatics resources frequently suffer from a lack of user-centered design (Helmy et al. 2016; Pavelin et al. 2012), and the trend could be related to this phenomenon.

**Table 2.** Updates Found for Currently Available Databases

| **Citation Quartile** | **Currently Available Count** | **Databases with Update Found Count (% within Quartile)** | **Update > 1 Year After Debut Count (% of found within Quartile)** |
| --- | --- | --- | --- |
| **1** | 155 | 111 (0.72) | 104 (0.94) |
| **2** | 232 | 137 (0.59) | 124 (0.91) |
| **3** | 236 | 114 (0.48) | 96 (0.84) |
| **4** | 193 | 72 (0.37) | 49 (0.68) |
| **Totals** | 816 | 434 | 373 |

Although a decreasing trend is also related to citation quartile (𝒳_2_ = 23.599, *p* = 1.186e-06), analysis of updates revealed that the majority of databases across all quartiles undergo updating after debut (Table 2). While this is helpful to confirm the databases are not static, the timing of updates is also informative as the recentness of update is indicative of continual maintenance. When analyzed, the most predominate update period across all quartiles was also the most recent and occurred in 2016-2017 (Figure 5). This result was significant for 1st (𝒳_2_ = 160.67, *p* < 2.2e-16), 2nd (𝒳_2_ = 108.73, *p* < 2.2e-16), and 3rd (𝒳_2_ = 25.386, *p* = 4.207e-05) quartiles, but not the 4th (𝒳_2_ = 5.6389, p = 0.2278).

**Figure 5.**
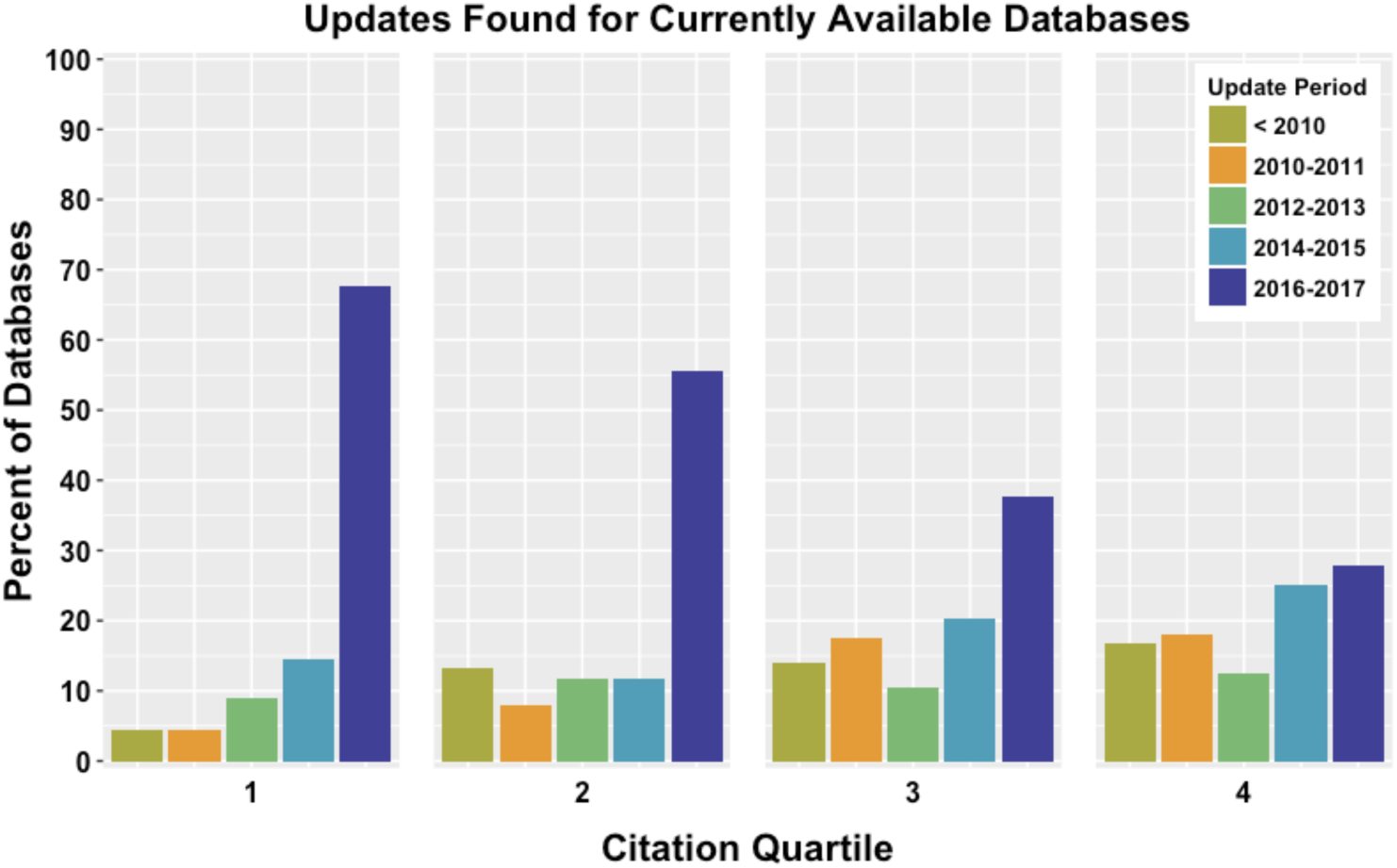
Database websites for currently available databases were checked for evidence of updating. Using the sample of databases binned into Citation Quartiles, databases with updates found were assigned to categories of 2-year increments. Updating is correlated to citation quartile, and the largest portion of databases examined were updated within the 2015-2017 time frame.

Especially bearing in mind this sample of databases was restricted to debut prior to 2013, these results show that for the majority of cases where documentation of updates could be found, databases are quite actively maintained. Recent updating was expected for databases that ranked more highly, but the results for the lower-ranking databases is somewhat more surprising. Although those in the 4^th^ quartile are not statistically likely to be updated in a given time period, that any update examples exist again provides notable outliers. This suggests that while citations are one measure on which to evaluate a database, other factors must be identified and explored to understand what motivates the on-going commitment to these resources.

## CONCLUSIONS

This work set out to gather a comprehensive census of molecular biology databases published in *NAR* Database Issues to provide a historical look at database development. The results here show rapid adoption of online molecular biology databases, with accumulation of over 1700 unique databases during the 25 year period covered. Moreover, new databases published within *NAR* Database Issues are proliferating at a rate of over 100 per year. This work reveals a tremendous appetite and capacity within the community to develop new resources.

Each of these databases was deemed of sufficient promise to be included in the prestigious *NAR* Database Issue. As such, each represents a significant amount of effort to bring it into existence. These efforts go well beyond acquiring hardware for computation and storage, or even that for development of the database’s core architecture, to also include a myriad activities around data acquisition and curation, quality control and validation, ingest workflows, interface design, and user support. With this in mind, the extent of proliferation uncovered here signals that the molecular biology and bioinformatics communities are remarkably energetic and willing to shoulder significant obligations in order to provide these resources to the boarder life science communities. During the initial stages of development it appears that funding, access to technology and skilled staff, and enthusiasm for the promise of such resources are not sufficiently limiting to prevent the extent of growth shown here. However, this study also shows that rapid growth of new resources is coupled with a rate of attrition that indicates the ability to establish new databases exceeds the ability to sustain those databases. While this reality was expected anecdotally, empirical evidence was lacking. As other domains likewise mature, a similar imbalance can be expected unless growth is more carefully metered or long-term support is more readily available.

In some regards, the results of the citation analyses presented here are not surprising. This work shows that the more cited a database, the more likely it is to be available and the more likely it is to have been updated recently. However, a result of these analyses is the identification of outlier populations which, if examined in more detail, have the potential to improve our understanding of the nuances related to database impact and persistence. For example, further analysis will help us understand where citation fails to anticipate availability, whether it’s for populations of high-ranking databases no longer available or populations of low-ranking databases somewhat unexpectedly available. Since considerable emphasis is currently being placed on increasing data citation, there is a danger that citations will disproportionately dominate arguments to justify either the continuation or discontinuation of support. This study shows that while citations and categorization into citation quartiles are useful as exploratory indicators, they are not conclusive in and of themselves. In keeping with the Leiden Manifesto on use of bibliometrics to evaluate research, it is critical that use of metrics must not result in “misplaced concreteness and false precision” (Hicks et al. 2015). Indeed, the outliers mentioned above appear especially well-placed to demonstrate the core of this principle, and the databases within this study are posed to serve as a prime example of the necessity of using metrics as a tool to assist expert qualitative evaluation instead of letting metrics stand as unquestioned judgements.

When evidence could be found, the analyses here shows that the majority of databases were updated recently. This held true even for many of the databases that mapped to the 3rd or 4th citation quartiles and indicates that most databases still available are also, in fact, actively maintained. As an initial evaluation, the criteria used here to determine what constituted maintenance was broadly defined, and the results suggest that a follow-up chronicling explicit maintenance activities would be both worthwhile and illuminative. In the meantime, this initial result is a highly positive finding for those who depend on reliable access to these resources as well as for those that argue the necessity of on-going, committed support. As funders, researchers, and database providers continue to grapple with sustainability issues, the ability to clearly articulate maintenance needs will become increasingly important. The work here provides the first step in this articulation by gathering preliminary evidence to show how frequently databases are updated in practice.

The updates observed here were reported in highly variable ways on database websites, and this serves as an area for improvement. Standardizing how activities are documented and reported would give anyone, including users, the ability to quickly assess the current status of a database’s operation. Some mechanisms, such as database certification, already exist and could facilitate the communication of such information. One such example is the recently debuted Core Trust Seal^3^, which merged the Data Seal of Approval and the World Data Systems certifications, is an inexpensive and light-weight process that is applicable to data resources across many domains. More specialized processes may also be appropriate, especially to capture domain-specific metrics. For example, organizations such as ELIXIR, an intergovernmental effort that coordinates and develops life science resources across Europe, is currently working to identify “Core Data Resources” and has outlined a number of indictors to evaluate these resources (Durinx et al. 2017).

Finally, as research becomes increasingly dependent on the availability of data through such community resources, we must be able to more nimbly address change. A consequence of the discrepancy between capacity to develop databases and the capacity to sustain them, particularly in light of increased public access to data, is the need to better embrace the many transitions a resource may undergo. While a database may not necessarily live in perpetuity as debuted, it need not dissolve into oblivion either. Within the databases studied here, numerous examples of proactive migration, merger, sunsetting, and archiving were witnessed. Indeed, database creation and database attrition seem to go hand-in-hand; an article that regretfully announced a discontinuation appeared in just the second Database Issue (Schmidtke and Cooper 1992). Yet these transitional activities are not widespread, let alone encouraged. Therefore, it is not surprising that examples of neglect, limbo, or complete abandonment were also found. As a community with a mature history of database development and use, promoting the legitimacy— and even necessity—of these proactive transitional activities would be a positive step forward. As other domains struggle with similar sustainability issues, a model that is more nuanced than “dead/alive” or “success/failure” would be valuable.

## SUPPORTING DATA

The data, scripts, and associated documentation for this article are archived and freely available in the Illinois Data Bank at https://doi.org/10.13012/B2IDB-4311325_V1. The author encourages corrections, reanalysis, reuse, and updating. Materials are also accessible on GitHub at https://github.com/1heidi/nar and may be updated since the time of this publication.

## ACKNOWLEDGEMENTS

The author would like to thank colleagues for helpful feedback on the manuscript, as well as the Illinois Statistics Office for their comprehensive methods review, Hoa Luong for careful curation of the dataset, and Colleen Fallaw for feedback on the manuscript and documentation as well as independent verification that the deposited R code executed as expected.

## Footnotes

1 “Online” used here refers to any networked access to resources on the Internet, including the current World Wide Web but also FTP or even earlier transfer protocols such as Gopher. “Web-available” used here is more specific and refers to availability through World Wide Web, which relies on HTTP, URLs, and browsers to provide access to resources on the Internet.

2 https://stats.datacite.org/

3 https://www.coretrustseal.org/

## REFERENCES

Agresti, Alan. 2007. An Introduction to Categorical Data Analysis. 2nd ed. New York: John Wiley & Sons.

Aphalo, Pedro J. 2016. Learn R… as You Learnt Your Mother Tongue. Leanpub. https://leanpub.com/learnr.

Attwood, Teresa K., Bora Agit, and Lynda B. M. Ellis. 2015. “Longevity of Biological Databases.” EMBnet.journal 21 (0): e803. https://doi.org/10.14806/ej.21.0.803.

Baker, Monya. 2012. “Databases Fight Funding Cuts.” Nature 489: 19. https://doi.org/10.1038/489019a.

Bastow, Ruth, and Sabina Leonelli. 2010. “Sustainable Digital Infrastructure: Although Databases and Other Online Resources Have Become a Central Tool for Biological Research, Their Long-term Support and Maintenance Is far from Secure.” EMBO Reports 11 (10): 730–34. https://doi.org/10.1038/embor.2010.145.

Baxevanis, Andreas D. 2000. “The Molecular Biology Database Collection: An Online Compilation of Relevant Database Resources.” Nucleic Acids Research 28 (1): 1–7. https://doi.org/10.1093/nar/28.1.1.

Dalgaard, Peter. 2008. Introductory Statistics with R. New York: Springer.

Durinx, Christine, Jo McEntyre, Ron Appel, Rolf Apweiler, Mary Barlow, Niklas Blomberg, Chuck Cook, et al. 2017. “sIdentifying ELIXIR Core Data Resources.” F1000Research 5 (March): 2422. https://doi.org/10.12688/f1000research.9656.2.

Ember, Carol, and Robert Hanisch. 2013. “Sustaining Domain Repositories for Digital Data: A White Paper.” In. http://datacommunity.icpsr.umich.edu/sites/default/files/WhitePaper_ICPSR_SDRDD_121113.pdf.

Fernández-Suárez, Xosé M., and Michael Y. Galperin. 2013. “The 2013 Nucleic Acids Research Database Issue and the Online Molecular Biology Database Collection.” Nucleic Acids Research 41 (D1): D1–7. https://doi.org/10.1093/nar/gks1297.

Galperin, Michael Y. 2006. “The Molecular Biology Database Collection: 2006 Update.” Nucleic Acids Research 34 (suppl_1): D3–5. https://doi.org/10.1093/nar/gkj162.

Galperin, Michael Y., and Guy R. Cochrane. 2009. “Nucleic Acids Research Annual Database Issue and the NAR Online Molecular Biology Database Collection in 2009.” Nucleic Acids Research 37 (suppl_1): D1–4. https://doi.org/10.1093/nar/gkn942.

Giannelli, Francesco, Peter M. Green, Katherine A. High, Steve Sommer, David P. Lillicrap, Michael Ludwig, Klaus Olek, et al. 1991. “Haemophilia B: Database of Point Mutations and Short Additions and Deletions—second Edition.” Nucleic Acids Research 19 (suppl): 2193–2220. https://doi.org/10.1093/nar/19.suppl.2193.

Gupta, Shashi, and Ram Reddy. 1991. “Compilation of Small RNA Sequences.” Nucleic Acids Research 19 (suppl): 2073–75. https://doi.org/10.1093/nar/19.suppl.2073.

Guthrie, Kevin, Rebecca Griffiths, and Nancy Maron. 2008. “Sustainability and Revenue Models for Online Academic Resources.” Ithaka. http://www.sr.ithaka.org/wp-content/uploads/2015/08/4.15.1.pdf.

Helmy, Mohamed, Alexander Crits-Christoph, and Gary D. Bader. 2016. “Ten Simple Rules for Developing Public Biological Databases.” PLOS Computational Biology 12 (11): e1005128. https://doi.org/10.1371/journal.pcbi.1005128.

Hicks, Diana, Paul Wouters, Ludo Waltman, Sarah de Rijcke, and Ismael Rafols. 2015. “Bibliometrics: The Leiden Manifesto for Research Metrics.” Nature News 520 (7548): 429. https://doi.org/10.1038/520429a.

Holdren, John P. 2013. “Increasing Access to the Results of Federally Funded Scientific Research.” Office of Science and Technology Policy. http://web.archive.org/web/20160115125401/https://www.whitehouse.gov/sites/default/files/microsites/ostp/ostp_public_access_memo_2013.pdf.

Imker, Heidi. 2018. “Molecular Biology Databases Published in Nucleic Acids Research between 1991-2016.” University of Illinois at Urbana-Champaign. https://doi.org/10.13012/B2IDB-4311325_V1

Jeanpierre, Cécile, Christophe Béroud, Patrick Niaudet, and Claudine Junien. 1998. “Software and Database for the Analysis of Mutations in the Human WT1 Gene.” Nucleic Acids Research 26 (1): 271–74. https://doi.org/10.1093/nar/26.1.271.

Jonkers, Koen, Gemma E. Derrick, Carmen Lopez-Illescas, and Peter Van den Besselaar. 2014. “Measuring the Scientific Impact of E-Research Infrastructures: A Citation Based Approach?” Scientometrics 101 (2): 1179–94. https://doi.org/10.1007/s11192-014-1411-

Kalumbi, Doyle, and Lynda B. M. Ellis. 1998. “The Demise of Public Data on the Web?” Special Features. Nature Biotechnology. December 1, 1998. https://www-nature-com.proxy2.library.illinois.edu/articles/nbt1298_1323.

Kirlew, Peter W. 2011. “Life Science Data Repositories in the Publications of Scientists and Librarians.” Issues in Science and Technology Librarianship 65. https://doi.org/10.5062/F4X63JT2.

Leydesdorff, Loet, Lutz Bornmann, Jordan A. Comins, and Staša Milojević. 2016. “Citations: Indicators of Quality? The Impact Fallacy.” Frontiers in Research Metrics and Analytics 1. https://doi.org/10.3389/frma.2016.00001.

MacRoberts, Michael H., and Barbara R. MacRoberts. 2017. “The Mismeasure of Science: Citation Analysis.” Journal of the Association for Information Science and Technology, n/a –n/a. https://doi.org/10.1002/asi.23970.

Marcial, Laura Haak, and Bradley M. Hemminger. 2010. “Scientific Data Repositories on the Web: An Initial Survey.” Journal of the American Society for Information Science and Technology 61 (10): 2029–48. https://doi.org/10.1002/asi.21339.

Martone, Maryann. (ed.). 2014. “Joint Declaration of Data Citation Principles - FINAL.” FORCE11. 2014. https://www.force11.org/datacitationprinciples.

Mayernik, Matthew S., David L. Hart, Keith E. Maull, and Nicholas M. Weber. 2017. “Assessing and Tracing the Outcomes and Impact of Research Infrastructures.” Journal of the Association for Information Science and Technology 68 (6): 1341–59. https://doi.org/10.1002/asi.23721.

Mayo, Christine, Todd J. Vision, and Elizabeth A. Hull. 2016. “The Location of the Citation: Changing Practices in How Publications Cite Original Data in the Dryad Digital Repository.” International Journal of Digital Curation 11 (1): 150. https://doi.org/10.2218/ijdc.v11i1.400.

Merali, Zeeya, and Jim Giles. 2005. “Databases in Peril.” Nature 435 (7045): 1010–11. https://doi.org/10.1038/4351010a.

Mooney, Hailey. 2011. “Citing Data Sources in the Social Sciences: Do Authors Do It?” Learned Publishing 24 (2): 99–108. https://doi.org/10.1087/20110204.

Neumann, Janna, Jan Brase, Janna Neumann, and Jan Brase. 2014. “DataCite and DOI Names for Research Data.” JOURNAL OF COMPUTER-AIDED MOLECULAR DESIGN 28 (10): 1035–41.

OECD. 2017. “Business Models for Sustainable Research Data Repositories,” December. https://doi.org/10.1787/302b12bb-en.

Pavelin, Katrina, Jennifer A. Cham, Paula de Matos, Cath Brooksbank, Graham Cameron, and Christoph Steinbeck. 2012. “Bioinformatics Meets User-Centred Design: A Perspective.” PLOS Computational Biology 8 (7): e1002554. https://doi.org/10.1371/journal.pcbi.1002554.

Schmidtke, Jörg, and David N. Cooper. 1992. “A Comprehensive List of Cloned Human DNA sequences—1991 Update.” Nucleic Acids Research 20 (suppl): 2181–98. https://doi.org/10.1093/nar/20.suppl.2181.

Southan, Christopher, and Graham Cameron. 2017. “D2.1: Database Provider Survey Report for ELIXIR Work Package 2.” Zenodo, May. https://doi.org/10.5281/zenodo.576013.

Tol, Paul. 2012. “Colour Schemes.” SRON/EPS/TN/09-002 Issue 2.2. SRON Netherlands Institute for Space Research. https://personal.sron.nl/∼pault/colourschemes.pdf.

Vines, Timothy H., Arianne Y. K. Albert, Rose L. Andrew, Florence Debarre, Dan G. Bock, Michelle T. Franklin, Kimberly J. Gilbert, Jean-Sebastien Moore, Sebastien Renaut, and Diana J. Rennison. 2014. “The Availability of Research Data Declines Rapidly with Article Age.” Current Biology 24 (1): 94–97. https://doi.org/10.1016/j.cub.2013.11.014.

Wada, Ken-nosuke, Yoshiko Wada, Hirofumi Doi, Fumie Ishibashi, Takashi Gojobori, and Toshimichi Ikemura. 1991. “Codon Usage Tabulated from the GenBank Genetic Sequence Data.” Nucleic Acids Research 19 (suppl): 1981–86. https://doi.org/10.1093/nar/19.suppl.1981.

Waltman, Ludo, and Michael Schreiber. 2013. “On the Calculation of Percentile-Based Bibliometric Indicators.” Journal of the American Society for Information Science and Technology 64 (2): 372–79. https://doi.org/10.1002/asi.22775.

Wells, Dan, and Doris Brown. 1991. “Histone and Histone Gene Compilation and Alignment Update.” Nucleic Acids Research 19 (suppl): 2173–88. https://doi.org/10.1093/nar/19.suppl.2173.

Wickham, Hadley. 2009. ggplot2: Elegant Graphics for Data Analysis. Springer-Verlag New York. http://ggplot2.org.

Wickham, Hadley. 2017a. Stringr: Simple, Consistent Wrappers for Common String Operations (version R package version 1.2.0). https://CRAN.R-project.org/package=stringr.

Wickham, Hadley 2017b. Tidyverse: Easily Install and Load “Tidyverse” Packages (version R package version 1.1.1). https://CRAN.R-project.org/package=tidyverse.

Wren, Jonathan D. 2016. “Bioinformatics Programs Are 31-Fold over-Represented among the Highest Impact Scientific Papers of the Past Two Decades.” Bioinformatics 32 (17): 2686–91. https://doi.org/10.1093/bioinformatics/btw284.

Wren, Jonathan D., Constantin Georgescu, Cory B. Giles, and Jason Hennessey. 2017. “Use It or Lose It: Citations Predict the Continued Online Availability of Published Bioinformatics Resources.” Nucleic Acids Research. https://doi.org/10.1093/nar/gkx182.

